# An energy costly architecture of neuromodulators for human brain evolution and cognition

**DOI:** 10.1101/2023.04.25.538209

**Authors:** Gabriel Castrillon, Samira Epp, Antonia Bose, Laura Fraticelli, André Hechler, Roman Belenya, Andreas Ranft, Igor Yakushev, Lukas Utz, Lalith Sundar, Josef P Rauschecker, Christine Preibisch, Katarzyna Kurcyus, Valentin Riedl

## Abstract

Humans spend more energy on the brain than any other species. However, the high energy demand cannot be fully explained by brain size scaling alone. We hypothesized that energy-demanding signaling strategies may have contributed to human cognitive development. We measured the energy distribution along signaling pathways using multimodal brain imaging and found that evolutionarily novel connections have up to 67% higher energetic costs of signaling than sensory-motor pathways. Additionally, histology, transcriptomic data, and molecular imaging independently reveal an upregulation of signaling at G-protein coupled receptors in energy-demanding regions. We found that neuromodulators are predominantly involved in complex cognition such as reading or memory processing. Our study suggests that the upregulation of neuromodulator activity, alongside increased brain size, is a crucial aspect of human brain evolution.

## Introduction

Over the course of 400 million years, the brains of various species have developed according to universal organizational principles (Clark et al., 2001; Sterling & Laughlin, 2017). Neurons are the local signaling units that form a dense connectome of widespread signaling pathways via their synapses. Notably, certain mammals have larger brain sizes, higher brain-to-body ratios, and greater neuron numbers than humans, and even Neanderthals had larger brains than homo sapiens (DeFelipe, 2011; Herculano-Houzel, 2012; Mortensen et al., 2014; Semendeferi et al., 2002; Smaers et al., 2021; Somel et al., 2013). This suggests that scaling the brain architecture is not sufficient for the emergence of complex cognition (Caceres et al., 2003; Fu et al., 2011; Preuss, 2011; Rakic, 2009; Roberts et al., 2022; Somel et al., 2013).

An alternative approach to uncover mechanisms related to human cognition is via its energy metabolism. The brain relies on a constant supply of energy substrates and, in humans, has the highest energy demand compared to the body (Heldstab et al., 2022; Martin, 1981; Navarrete et al., 2011; Pontzer et al., 2016). This suggests that energy should be distributed efficiently and according to its value for information processing (Bullmore & Sporns, 2012; Conrad et al., 2017; Laughlin, 2003; Laughlin et al., 1998). On a cellular level, the energetic costs of signaling of individual neurons have been optimized across evolution and are stable across different mammals (J. J. Harris et al., 2015; Herculano-Houzel, 2011; Howarth et al., 2012; Hyder et al., 2013; Karbowski, 2012; Laughlin, 2001). On a systems level, the energy demand of larger functional systems is, however, unknown. We hypothesized that the distribution of energy metabolism along signaling pathways will reveal mechanisms of complex information processing in the human brain.

## Results

We quantified the energetic costs of signaling as the relationship between cortical glucose metabolism and synchronized signaling across the brain connectome. Healthy subjects were scanned on an integrated PET/MRI-scanner to simultaneously measure the cerebral metabolic rate of glucose (CMRglc) and the extent of synchronized signaling between brain voxels, i.e., the degree of functional connectivity (dFC) (Fig. 1A).

**Fig. 1:**
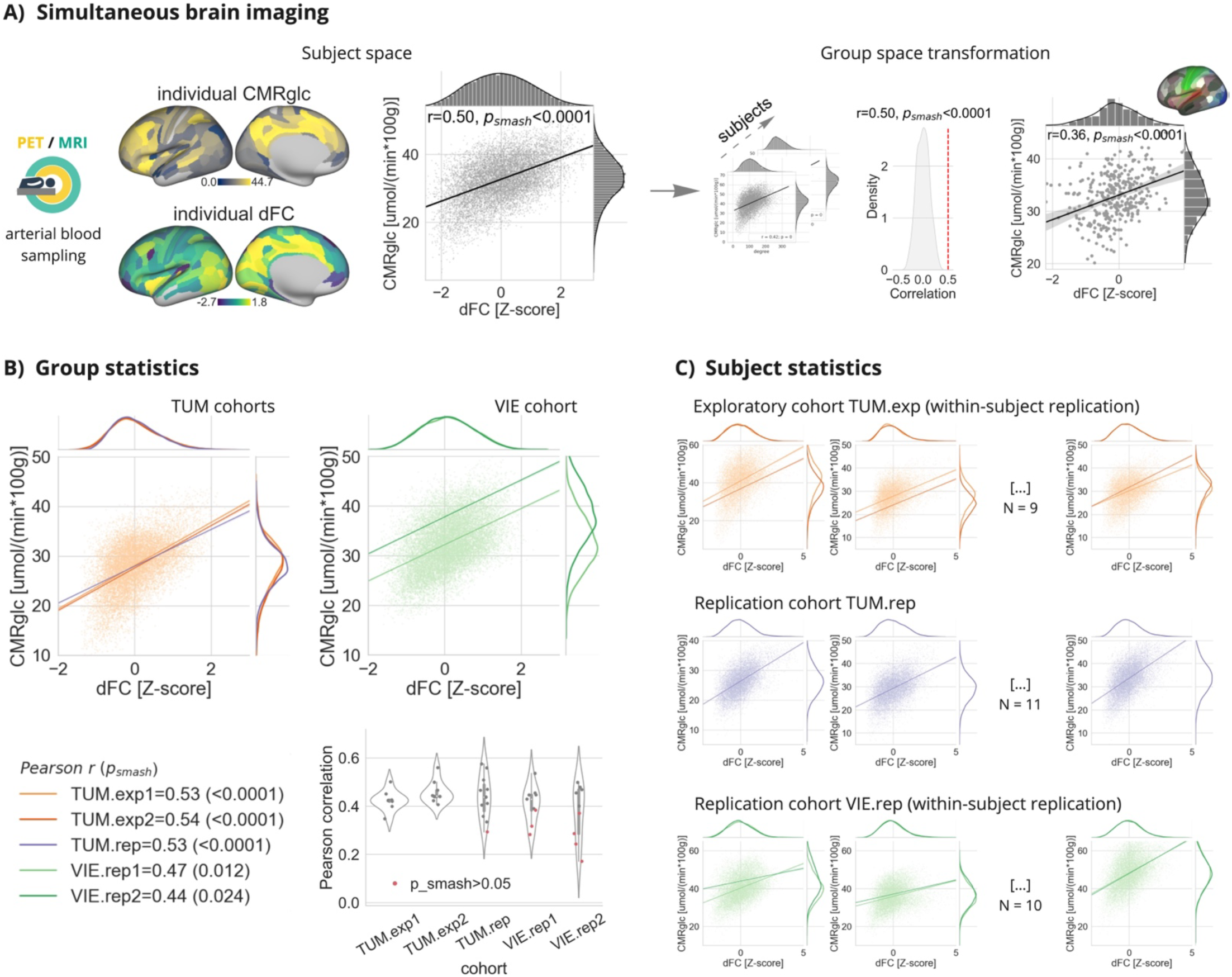
Glucose metabolism scales linearly with the degree of functional connectivity in individual brains. (**A**) We analyzed the voxel-wise relationship of CMRglc and dFC in the individual brain space of each subject from an exploratory cohort measured twice at our institution (TUM.exp1/2: light/dark orange), from a younger replication cohort measured again at our institution (TUM.rep: violet), and from a second replication cohort of healthy subjects that were scanned twice at a different institution (VIE.rep1/2: light/dark green). For group statistics, brain data were transformed into a standard brain space (MNI, see Methods) on either the level of voxels or averaged for functional regions of a standard parcellation atlas (MMP, see Methods). (**B**) Significant Pearson’s correlation between voxel-wise CMRglc and dFC in MNI space averaged across individuals from each of the five groups. The distribution of Pearson’s *r*-values of each individual dataset are summarized in violin plots. (**C**) Exemplary plots of significant Pearson’s correlation between voxel-wise CMRglc and dFC of individual datasets. Regression plots of all subjects can be found in Fig. S1 of the Supplementary material.

We identified a linear relationship between CMRglc and dFC in each individual brain of three different cohorts that were measured at two different institutions (Fig. 1B, C). We first analyzed data from an exploratory cohort (TUM.exp1 and TUM.exp2, age: 43 ± 7 years, 4 females, *N* = 9, measured twice) and replicated our results in three other datasets from two cohorts (TUM.rep cohort, age: 27 ± 5 years, 6 females, *N* = 11, measured once; VIE.rep1 and VIE.rep2 cohort, age: 27 ± 7 years, 5 females, *N* = 10, measured twice) using the identical analysis pipeline (Fig. 1B; all *p_smash* < 0.024, correcting for spatial autocorrelation with permutation testing, see Methods). Fig. 1C shows exemplarily plots of individual subject data, while the results of each subject and imaging session can be found in Fig. S1 of the Supplemental Material (Pearson’s *r*: mean = 0.42, s.d. = 0.08; all *p* < 0.1 * 10^-24, from individual correlation analyses, *N* = 47; *p_smash* > 0.05 for *N* = 8 datasets).

There was no significant difference in Pearson’s *r* between cohorts (Pearson’s *r* range = [0.17, 0.57]; mean = 0.42; s.d. = 0.08; *F*(1,28) = 2.41, *p* = 0.13, one-way ANOVA). Please note that across cohorts and scanners, we identified a similar increase in energy demand per connection (slope range = 1.12 - 3.79; mean = 2.42; s.d. = 0.69; *F*(1,28) = 0.07, *p* = 0.79, one-way ANOVA), but varying baseline CMRglc (y-intercept). Furthermore, the linear increase in energetic costs is independent of sex and age (Pearson’s *r* female / male: mean = 0.44 / 0.41, standard deviation (s.d.) = 0.10 / 0.05; *F*(1,28) = 0.82, *p* = 0.37, one-way ANOVA; Pearson’s *r* age = 0.07; *p* = 0.70; CI: [-0.30, 0.42]; *N* = 30 subjects). We also found no effect of gray matter partial volume on the energy-connectivity scaling using voxel-based morphometry as an explanatory variable in the linear model (*F*(2,330) = 50.05, *p* = 0.13, one-way ANOVA). Finally, we tested the stability of the energy-connectivity scaling for different connectivity measures. Results show a consistent scaling of energetic costs of signaling with both dynamic functional and structural connectivity (Fig. S2 of the Supplementary Materials). We deposited scripts and data in online repositories, and one can replicate individual steps of the analysis in an online notebook (see Data and code availability).

Next, we identified regions with deviating energetic costs of signaling calculated as the residual CMRglc per dFC (Fig. 2A, upper row). Across all cohorts and individual brains, we identified a consistent cortical distribution of energetic costs for signaling. The brain map in Fig. 2A shows regions of higher (red) and lower (blue) energetic costs averaged across subjects of the cohort TUM.exp1. We performed identical analyses on the remaining four datasets and found a consistent cortical energy distribution in each of the cohorts, with higher energetic costs of signaling in frontal and lower costs in sensory cortices (Fig. 2B; significant spatial similarity between the pattern of each cohort and that of TUM.exp1, all *p_smash* < 0.0001, voxel-wise permutation test (two-sided) preserving spatial autocorrelation, 1000 permutations). Fig. 2C shows the average map of energetic costs of signaling average across all subjects of all cohorts. Please note that residual CMRglc varies by +/- 25% across the connectome compared to an average metabolic rate of CMRglc = 31.4 µmol / (min * 100 g).

**Fig 2:**
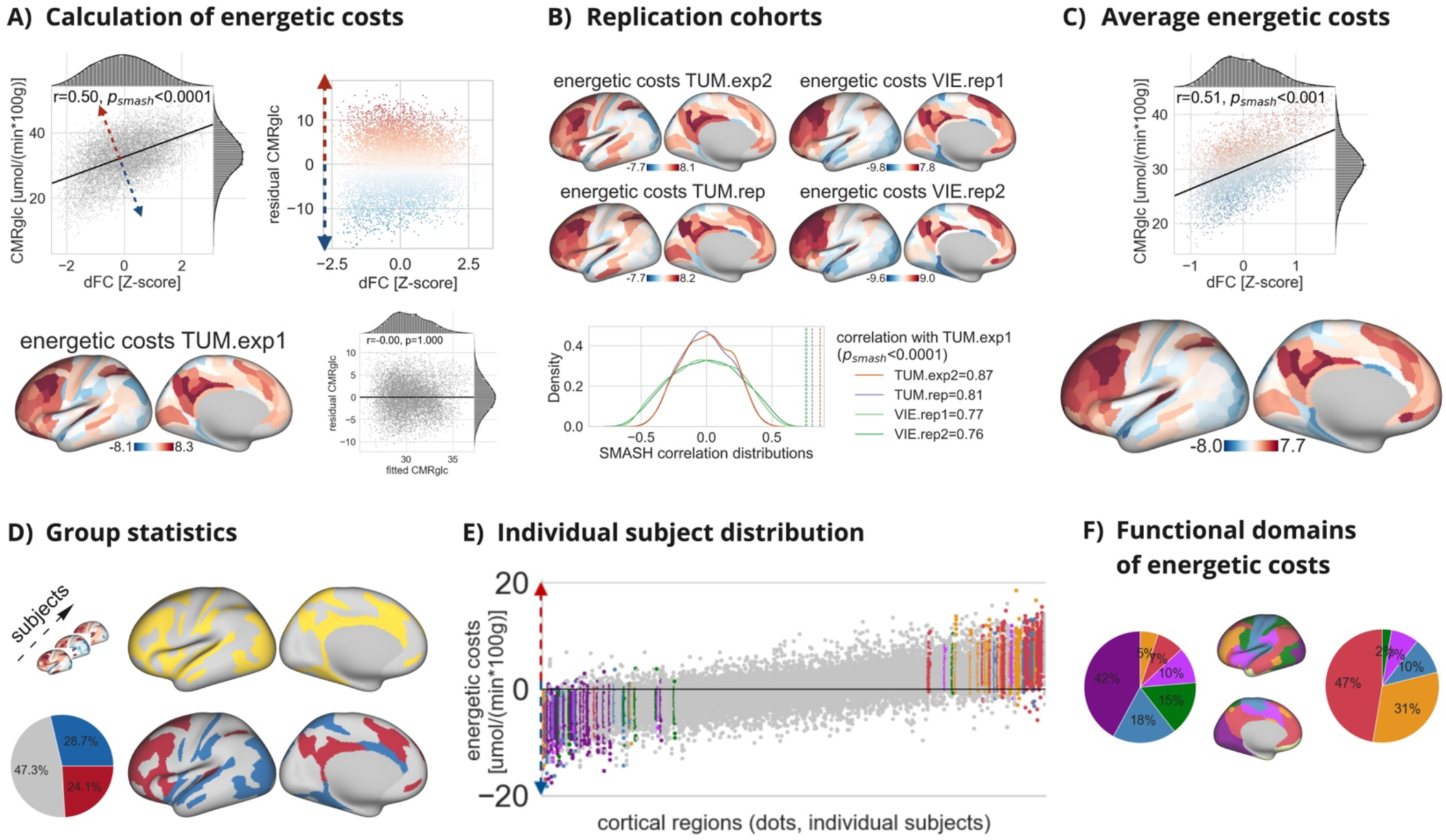
Regional distribution of energetic costs of signaling across the brain connectome. (**A**) Upper row: Scatterplot of dFC (x-axis) and CMRglc (y-axis) for a single subject with corresponding distribution of residual CMRglc showing voxels with higher (red) and lower (blue) energy demand per dFC. Lower row: Brain map of average energetic costs of signaling in cohort TUM.exp1. The scatterplot of fitted (x-axis) vs. residual (y-axis) CMRglc across all subjects shows a random distribution, indicating no unexplained structure in the model (*r* = 0, *p* = 1, CI = [-0.03, 0.03]). (**B**) Significant spatial similarity between the distribution of energetic costs of signaling in TUM.exp1 and each of the four cohorts. (**C**) Voxel-wise scatter plot and region-wise brain surface of energetic costs of signaling averaged across the subjects of all cohorts. (**D**) Brain surface showing regions with significantly deviating (yellow), and specifically with higher (red) and lower (blue) energetic costs of signaling. Pie chart summarizes the distribution of regions with varying energetic costs. (**E**) Strip plot illustrates the region-wise (x-axis) distribution of energetic costs of signaling for each subject of all cohorts (dots) sorted from low-to-high median costs. Regions are colored if energetic costs deviate from the normalized cortex mean in > 95% of all subjects. Color codes illustrate the affiliation of regions to one of seven normative functional networks (see Methods). (**F**) Pie charts summarize the affiliation of regions with lower (left) and higher (right) energetic costs of signaling to any of the seven normative brain networks projected on the brain surface.

We then investigated whether energetic costs of signaling relate to certain functional domains of the cortex. Fig. 2D shows the result of a group analysis identifying cortical regions with significant deviations in CMRglc against the null hypothesis of the linear model fit (yellow; *p* < 0.01, voxel-wise non-parametric permutation t-test (two-sided) with 5000 permutations, corrected for multiple comparisons using the family-wise error rate). Results show that 28.7% of all regions have significantly lower (blue) and 24.1% of regions have significantly higher (red) energetic costs than predicted by the model. On the subject level, we identified a set of 76 regions (23% of all regions) with diverging energetic costs in > 95% of all datasets (colored ROIs in Fig. 2E; dots represent single subjects) and a core set of 23 regions (7% of all functional regions) with deviating energetic costs in each subject of all datasets. The color scheme in Fig. 2F indicates the affiliation of cost-deviating regions to one of seven established functional networks (see Methods). The pie charts show that regions with lower energetic costs cluster in sensory-motor networks (violet, blue, and green sum up to 75%), while regions with higher energetic costs are mainly located in fronto-parietal networks (red and yellow sum up to 78%).

In summary, we identify a consistent pattern of varying energetic costs of signaling pathways in individual subject data. In particular, the energy demand for signaling in fronto-parietal networks is up to 67% higher than in sensory-motor networks.

Given the higher energetic costs of certain signaling pathways, we next investigated whether the human brain has an overall higher glucose metabolism as compared to other species. The biological scaling of metabolism in relation to the size or volume of an organ is called metabolic allometry. We replicated the scaling exponent of 0.86 (Karbowski, 2007) for the log – log relationship between cortical glucose metabolism and brain volume across ten mammals (least-square fit, *log(glc)* = *log(volume)*^0.86 - 0.09; *R*^2^ = 0.993; *p* < 0.0001; *N* = 10). The average glucose metabolism in our human cohorts (CMRglc = 31.35 µmol / (min * 100g)) lies within the confidence interval of the least-square fit (Fig. 3A; CI: [0.81, 0.91]). This means that the energy metabolism of the entire human brain is as high as predicted for its size by the law of metabolic allometry.

**Fig 3:**
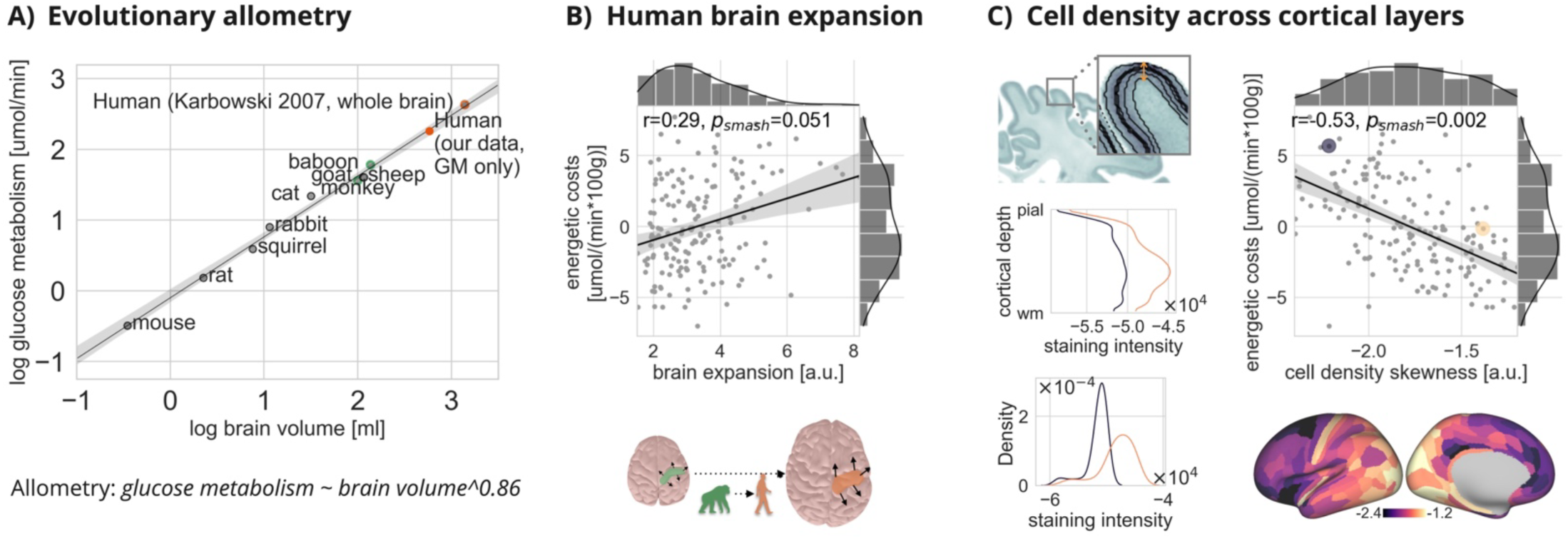
Energetic costs of signaling relate to the evolution of brain morphology on the macro- and microscale. (**A**) Allometric scaling of brain glucose metabolism with volume across ten mammals including humans (external data, see Methods). Our data from grey matter (GM) fall within the confidence interval of the model fit (orange dot). (**B**) Significant Pearson’s correlation between energetic costs of signaling and the degree of cortical expansion from non-human to human primates. (**C**) Energetic costs of signaling correlate with cell density in infragranular layers. Histological slice of the BigBrain atlas (see Methods) showing the staining intensity for cells in the gray matter (**top**). The distribution of staining intensity for cells between pial and white matter (WM) surfaces is shown for two exemplary regions (violet, yellow) which is then translated into the skewness of staining intensity (**bottom**). The two regions are examples for high cell density in supragranular (yellow, right tailed) and infragranular (violet, left tailed) layers. The brain surface shows the cortical distribution of layer predominance across the entire cortex. The scatterplot reveals a significant negative Pearson’s correlation between energetic costs of signaling and cell density skewness, indicating highest energetic costs in regions with highest cell density in infragranular layers.

Are energy demanding regions specific to human evolution? Using a morphometric atlas that describes the expansion of homologous brain regions from chimpanzees to humans (see Methods), we found a positive linear relationship between the energetic costs of signaling and brain expansion (Fig. 3B; *r* = 0.29, *p* < 0.0001; *p_smash* = 0.051, CI: [0.42, 1.04]; *N* = 168 regions from MMP-parcellation). Please note that the x-axis indicates the extent of expansion during evolution and not the actual size of a region. In summary, the entire human brain evolved according to the allometric scaling law of metabolism. However, regions that expanded most during human evolution show an excessive energy demand compared to the rest of the brain.

In a next step, we studied the microstructure of brain regions with high energetic costs of signaling by means of the BigBrain Atlas (see Methods). This is a histological atlas of the cellular distribution across 50 cortical layers of a human donor brain (Fig. 3C). For each region, the staining intensity of brain cells is plotted along the cortical depth and transformed into a density distribution. The density skewness indicates the relative distribution of cells along the cortical depth (*left skewness* ∼ *higher cell density in lower layers*). We then calculated the correlation between energetic costs of signaling and the cell density distribution across cortical regions. Results show a significant negative relationship between energetic costs and the skewness towards upper layers (Fig. 3C, right; *r* = -0.53, *p* < 0.0001, *p_smash* = 0.002, CI: [-6.89, -4.53]; *N* = 168 MMP-regions). This means that regions with high energetic costs of signaling have a higher cell density in lower cortical layers compared to regions with low energetic costs.

Given the difference in macroscale and microscale morphology, we hypothesized a unique molecular profile associated with the distribution of energetic costs of signaling. The Allen Human Brain Atlas (AHBA) provides transcriptomic data sampled across the cortical surface and averaged across six donor brains (see Methods). We projected the AHBA microarray data onto our map of energetic costs of signaling and performed pairwise correlations between regional costs and each of the expression profiles of 8426 brain specific genes across cortical regions. Results show that 617 gene expression profiles significantly correlate with energetic costs of signaling across the human cortex (Fig. 4A, left; *p* < 0.005, FDR-corrected for multiple comparisons). We then investigate the putative function of the significantly correlated genes using a gene ontology enrichment (GOE) analysis. The analysis of ‘cellular components’ revealed that regions with high energetic costs of signaling are significantly enriched in genes coding for synapses, part of synapses and dendrites, i.e. cellular compartments that are involved in signal transduction (Fig. 4A, middle; *p* < 0.02, FDR-corrected; Table S1). The analysis of ‘molecular functions’ identified seven significantly enriched clusters, also predominantly related to signal transduction (Fig. 4A, right; *p* < 0.05, FDR-corrected; Table S2). Specifically, we found gene annotations for activity of receptors and transporters involved in metabotropic, i.e. G-protein-coupled, neuromodulation and voltage-gated signaling. Results were replicated using a different GOE-tool and database (see Methods, Tables S3, S4). A summary of the ‘molecular functions’ is depicted in the stacked pie chart of Fig. 4B. This shows that 95% of genes that are overexpressed in regions with high energetic costs are involved in signal transduction (green), and mainly in metabotropic signaling (pink, 40%). In other terms, the human brain spends excessive energy on the long-lasting regulation of (fast) neurotransmission with (slow) neuromodulators such as serotonin, dopamine or noradrenaline.

**Fig 4:**
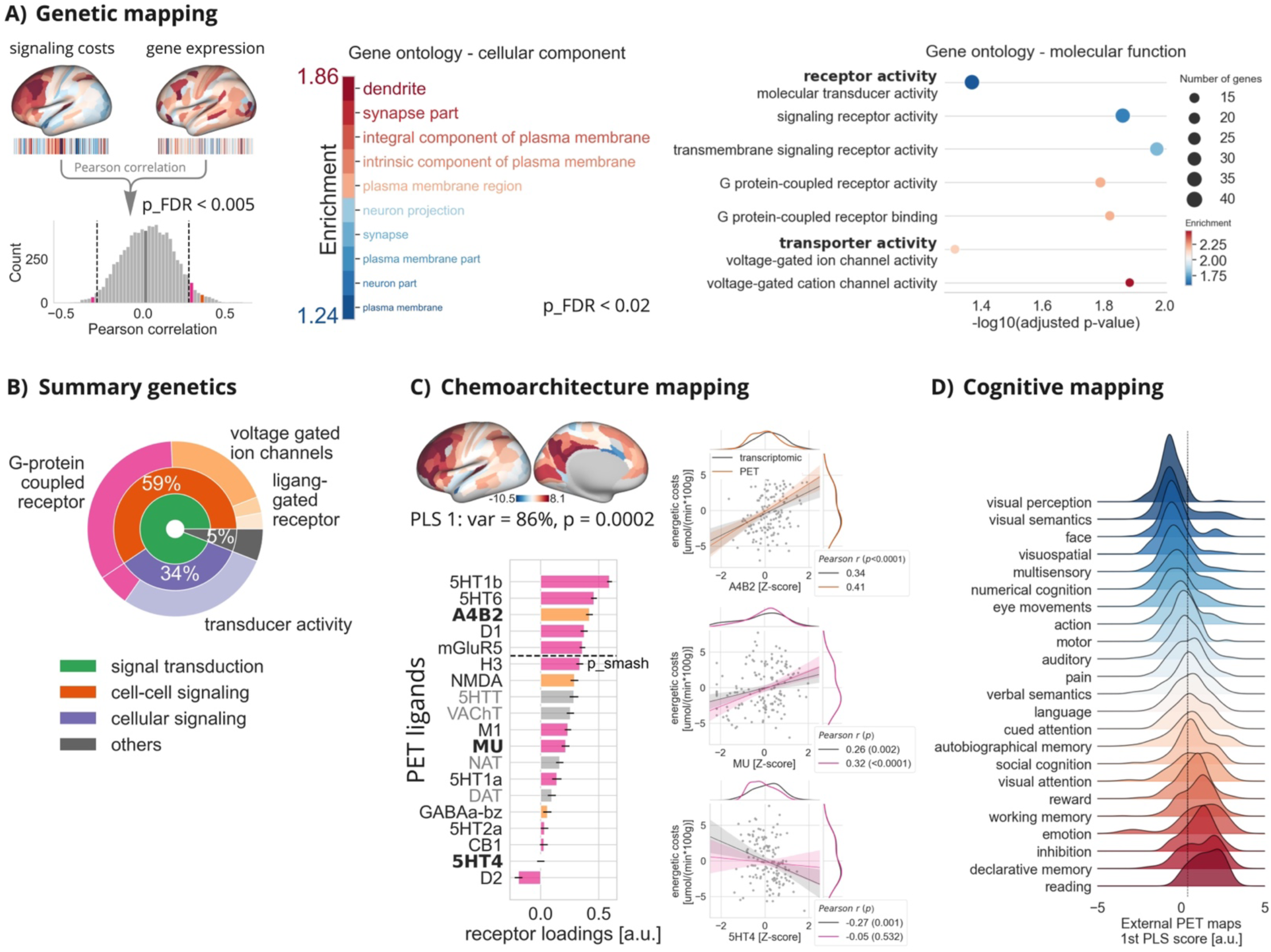
Regional activity of neuromodulators relates to high energetic costs of signaling and complex cognition. (**A**) Significant Pearson’s correlations (dashed lines) between energetic costs and each of 8.426 brain specific gene expression values across 170 cortical regions. Gene ontology enrichment analysis identifies cellular components (**middle**) and molecular functions (**right**) that are significantly associated with correlated genes. (**B**) Hierarchical summary of the gene ontology enrichment analysis stressing the involvement of genes coding for proteins involved in signal transduction (95%, green), and particularly of G-protein coupled, metabotropic, neuromodulation (40%, pink) and ionotropic signaling (26%, orange). (**C**) Multi-dimensional PLS-analysis reveals that 86% of the cortical distribution of energetic costs (first latent variable, PLS 1) is explained by the linear combination of neurochemical signaling as defined by receptor-PET imaging of 19 neuromodulators and neurotransmitters (metabotropic: pink, ionotropic: orange, transporters: grey). Error bars represent 95% confidence intervals using bootstrap resampling (5000 permutations). (**Right**) For three receptors, both transcriptomic (colored bars in **A**) and imaging (’bold’ ligands in **C**) data are available which allows to validate activity of unique receptors with gene expression data. For the ionotropic (A4B2) and the two metabotropic (MU, 5HT4) receptors, the separate regression analyses between energetic costs and both transcriptomic and imaging data yield similar directions and slopes (colored regressions: PET imaging, grey regressions: transcriptomic data), supporting the results from imaging with transcriptomic data. (**D**) Joyplot shows histograms of the voxel-wise first latent score between energetic costs and chemoarchitecture profiles (z-score > 2.3) for each of 23 cognitive domains (y-axis) from the meta-analytic Neurosynth database (see Methods). Color-coding illustrates the increasing rank (blue > red) of average energetic costs from simple sensory processing to higher cognitive functions.

Next, we replicated this finding about the genetic upregulation of neuromodulator activity with receptor imaging data. In a strong collaborative approach, colleagues recently gathered PET imaging data about the brain-wide activity of 19 unique receptors and transporters from 28 different PET studies (see Methods). This includes receptor-activity maps of the serotonergic (5HT1b, 5HT6, 5HT1a, 5HTT, 5HT2a, 5HT4), dopaminergic (D1, FDOPA, DAT, D2), cholinergic (A4B2, VAChT, M1), adrenergic (NAT), histaminergic (H3), cannabinoid (CB1), and opioid (MU) system, in addition to three receptors of the glutamatergic (mGluR5, NMDA) and GABA-ergic (GABAa-bz) system. We entered all PET maps of ligand occupancy into a partial least-squares regression (PLS) analysis to test the joint explanatory power of neuromodulator activity for the spatial distribution of energetic costs of signaling. The first component of the PLS analysis (brain map in Fig. 4C) explains 86% of the cortical variance in energetic costs (*p* = 0.0002 after 5000 randomizations). Specifically, 16 receptors and transporters significantly contribute to the main latent variable (bar plots in Fig. 4C). This means that the cortical distribution of energetic costs of signaling is strongly related to the regional level of neuromodulator activity.

We finally validated the consistency between transcriptomic- and PET imaging data for individual receptors. Three of the 617 overexpressed genes from the AHBA specifically code for membrane receptors whose density was captured by unique PET-ligands respectively (see ‘bold’ ligands in Fig. 4C; receptor / gene / PET-ligand combination listed here: opioid receptor mu / OPRM1 / [11 C]-carfentanil; serotonergic receptor 5HT4 / HTR4 / [11 C]-SB207145; cholinergic receptor alpha-4 beta-2 / CHRNA4 / [18 F]-flubatine). This allowed us to specifically test the consistency between gene expression levels and imaging-based activity for unique receptors. For each of the three receptors, ligand-activity confirmed the level of gene expression with similar signs and slopes of regressions across cortical regions (Fig. 4C, right; PET data: MU: *r* = 0.32, *p* < 0.0001, *CI* = [0.17, 0.46]; 5HT4: *r* = -0.05, *p* = 0.532, CI: [-0.21, 0.11]; A4B2: *r* = 0.41, *p* < 0.0001, CI: [0.26, 0.53]). Together, transcriptomics and molecular imaging independently suggest a high level of neuromodulator activity in energetically expensive regions of the brain. Particularly, our analyses point towards excessive energy demands for long-lasting, G-protein coupled neuromodulation.

So far, we have identified a high density of slow-acting neuromodulator activity in evolutionarily novel cortex. In a final step, we explored whether these regions are involved in higher cognitive processing. This would support the notion of an expensive signaling architecture being dedicated to human cognition. The Neurosynth project is a meta-analytic database with statistical maps aggregating voxel-wise activity for a wide range of cognitive functions derived from thousands of neuroimaging studies (see Methods). We extracted the regional activity maps for 23 cognitive domains ranging from simple sensory processing to complex cognition and evaluated the similarity between the chemoarchitecture map of Fig. 4C with each activity map. Results show that regions with strong neuromodulator activity particularly contribute to complex functions such as memory processing and reading but are less prominent in activity patterns of sensory-motor processing (Fig. 4D).

## Discussion

Using a unique imaging setup, we quantified the energetic costs of signaling across the human brain. We identified an excessive demand for the long-term regulation of neurotransmitter signaling via neuromodulators. Intriguingly, neuromodulator activity is dense in the evolutionarily novel cortex and is particularly involved in cognitive processing.

On a global level, we confirm that total cerebral energy metabolism scales according to the law of metabolic allometry (Balaban, 2013; Karbowski, 2007; West et al., 2002). This means that the human brain consumes less energy per volume than its predecessors. It has been suggested that this is due to an energy-efficient global signaling architecture in humans (Bullmore & Sporns, 2012; Conrad et al., 2017; Laughlin, 2001; Preuss, 2011; Sterling & Laughlin, 2017). We measured an average metabolic rate of 31.35 µmol glucose/min per 100g of gray matter tissue, which is the equivalent of around 12 cubes of sugar that are metabolized by an average-sized human brain per day. On the regional level, however, we identified up to 50% variance in energetic costs for individual signaling pathways. Specifically, we noticed higher energy metabolism in evolutionarily novel structures. This regional variance in metabolism corresponds with findings from transcriptomic and metabolomic analyses of post-mortem brains from humans and primates, which suggest metabolic upregulation in the frontal cortices (Caceres et al., 2003; Fu et al., 2011; Somel et al., 2013; Wei et al., 2019).

What is the mechanism behind the excessive energy demands in evolutionarily novel cortex? Using transcriptomic and receptor imaging data, we find that excessive energy metabolism in evolutionarily novel cortex is related to neuronal signaling and particularly to neuromodulator activity. These neuromodulators, such as serotonin, dopamine, and noradrenaline, act as general modulators of brain-wide circuits (Grossman & Cohen, 2022; Nadim & Bucher, 2014). The G-protein-mediated regulation of fast neurotransmitter signaling creates a long-lasting and widespread effect on information processing. This effect is more about setting the tone of general excitability than transferring individual bits of information.

The greater energy demand of slow-acting neuromodulators compared to fast neurotransmitter signaling is due to a sequence of biochemical steps that include second messengers and protein transformations (Greengard, 2001). This confirms our observation of excessive energy demand for regions with a large number of neuromodulators in addition to a linear relationship between energy expenditure and the degree of signaling pathways. Regions with an upregulation of metabotropic signaling also have a distinct cellular and genetic composition. Our analysis of histological data indicates a higher cell density in lower cortical layers, while transcriptomic data show an enrichment of genes associated with signal integration at dendrites and synapses. This is supported by recent autoradiography data that links information integration to cellular components, particularly those in lower cortical layers (Goulas et al., 2021).

Recent research has suggested that the energetic costs of signaling increase during long-term processes such as memory formation (Bazzari & Parri, 2019). Our human data indicate that brain regions with high energetic costs of metabotropic signaling play an important role in cognitive processing over longer timescales, including memory processing, cognitive inhibition, and reading. Additionally, neuromodulators have been linked to cognitive dysfunctions associated with major mental disorders (Roth, 2019). Unfortunately, the efficacy of current psychoactive drugs in regulating neuromodulators is limited, and further research is needed to better understand the dysfunction of metabotropic signaling in those patients.

Our findings suggest that the evolution of human cognition may have emerged not only from an overall larger brain, but particularly by the development of slow-acting neuromodulator circuits. It seems that the benefits of increased cortical energy metabolism, together with an increased supply of energy substrates (Heldstab et al., 2022; Navarrete et al., 2011; Pontzer et al., 2016), have outweighed its risks. Yet, our knowledge of how the interaction of slow-acting neuromodulators with fast information processing contributes to human cognition is still limited.

## Acknowledgments

Valentin Riedl is supported by the European Research Council (ERC) under the European Union’s Horizon 2020 research and innovation programme (ERC Starting Grant, ID 759659). We thank Pawel Markiewicz for his technical support in the use of the NiftyPET library; Thomas Beyer for sharing the PET/MRI dataset from the VIE-site; Martjin van den Heuvel and Yongbin Wei for their help in using the cortical expansion from non-human primates to humans dataset; and Markus Ploner for his feedback about the manuscript. We thank our colleagues at the Technical University of Munich, particularly Claus Zimmer, Wolfgang Weber, Gerhard Schneider, Sylvia Schachoff, and Jorge Cabello.

## Data and code availability

All raw and processed data from the TUM-site are available in the online repository of OpenNeuro(Castrillon et al., 2023) (https://openneuro.org/datasets/ds004513), whereas the data from VIE-site may be requested from the co-author (L.S.). The scripts, python jupyter notebooks, and configuration files for replication of all analyses and figures are available on github (https://github.com/NeuroenergeticsLab/energetic_costs), as is a link to a ready-to-run, interactive notebook to generate all results from this manuscript via binder.

## Author contributions

V.R.: conceptualization and supervision. G.C.: methodology and validation. G.C. and V.R.: visualization and writing of the original draft. G.C. and L.S.: data curation. A.B., A.R., C.P., I.Y., J.R., K.K., L.S., L.U. and S.E.: investigation. A.H., G.C., L.F., L.S., L.U. and R.B.: formal Analysis. All authors: writing, review and editing of the draft.

## Supplemental information

Figs. S1 to S2 Tables S1 to S4

## Materials and Methods

The data processing was performed using the Python packages Pandas (*Pandas - Python Data Analysis Library*, n.d.) and Numpy (C. R. Harris et al., 2020), the neuroimaging data were handled using Nilearn (Abraham et al., 2014) and Nibabel (Brett et al., 2022), the plotting was performed with Matplotlib (Hunter, 2007), Seaborn (Waskom, 2021) and Joyplot (Taccari, 2017/2023); whereas the brain surface representations were printed using WBplot (J. Burt, 2020/2022).

### Participants

Forty-seven datasets from thirty healthy participants from three independent cohorts were included in the study. Three additionally available datasets had to be excluded, two due to motion artifacts with framewise displacement > 0.25 mm (Power et al., 2014) and one due to incomplete data. All participants were right-handed and did not report any history of psychiatric conditions. Participants were informed about the objectives and potential risks of the study, and signed a written consent inform. The study was approved by the local institutional review board and was conducted in accordance with the Declaration of Helsinki. Three cohorts of participants were analyzed, two of them were recruited from our site and another from an external site: i) a within-subject exploration sample (TUM.exp) of 9 participants (mean age = 43 years, std = 7 years; 4 females); ii) a prospective replication sample (TUM.rep) of 11 participants (mean age = 27 years, std = 5 years; 6 females); and iii) an external within-subject replication sample (VIE.rep) (Sundar et al., 2018) of 10 participants (mean age = 27 years, std= 7 years; 5 females). Two participants from the VIE.rep cohort had only one session.

### Data acquisition

At TUM, we simultaneously measured FDG-PET activity and BOLD-fMRI signals during resting conditions while the participants kept their eyes open, except for the second imaging session of TUM.exp, where the participants had their eyes closed. Data were acquired on an integrated PET/MR (3T) Siemens Biograph mMR scanner (Siemens, Erlangen, Germany) and used a 12-channel phase-array head coil for the MRI acquisition. The PET data were collected in list-mode format with an average intravenous bolus injection of 184 MBq (s.d. = 12 MBq) of [18F]FDG. In parallel to the PET measurement, automatic arterial blood samples were taken from the radial artery every second to measure blood radioactivity using a Twilite blood sampler (Swisstrace, Zurich, Switzerland).

The functional MRI data were acquired during a 10 min time interval using a single-shot echo planar imaging sequence (300 volumes; 35 slices; repetition time, TR = 2000 ms; echo time, TE = 30 ms; flip angle, FA = 90°; field of view, FOV = 192 × 192 mm^2^; matrix size = 64 × 64; voxel size = 3 × 3 × 3.6 mm^3^). Diffusion-weighted images were acquired using a single-shot echo planar imaging sequence (60 slices; 30 non-colinear gradient directions; b-value = 800 s/mm² and one b=0 s/mm² image; TR = 10800 ms, TE = 82 ms; FA = 90°; FOV = 260 x 264 mm²; matrix size = 130 x 132; voxel size = 2 x 2 x 2 mm^3^). Anatomical images were based on a T1-weighted 3D-MPRAGE sequence (256 slices; TR = 2300 ms; TE = 2.98 ms; FA = 9°; FOV = 256 × 240 mm^2^; matrix size = 256 × 240; voxel size = 1 × 1 × 1 mm^3^).

The acquisition and formats of data from the VIE-site were similar to the TUM-site and described elsewhere (Sundar et al., 2018). The main differences were a higher dose and variability of injected [18F]FDG (mean dose = 356 MBq, std = 66 MBq), manual measurement of the blood radioactivity for the arterial input function, and a slightly different fMRI protocol with acquisition time (∼7 min), TR (2400 ms) and voxel size (2 × 2 × 3.7 mm^3^).

### Data processing

The following pre- and postprocessing pipelines for MR- and PET-data were established on the TUM.exp-dataset and applied without further modification to the TUM.rep- and VIE.rep-datasets, except for reasons of different data structure between TUM- and VIE-data which is explicitly stated below. All processing steps and calculation of parameter maps were performed in native space of each modality and only transformed for group analyses into a standard brain space of the Montreal neurological institute MNI152NLin6ASym with 3 mm voxel resolution. For the region-of-interest (ROI) analyses, we used the HCP-MMP1.0 parcellation scheme, a population-based cortical parcellation with 180 ROIs per hemisphere from the Human Connectome Project dataset (Glasser et al., 2016) and calculated the median value of the metric of interest per ROI. Furthermore, each ROI was labeled according to its location within one of seven normative brain functional networks (Yeo et al., 2011).

#### MRI

The preprocessing of the structural and functional MRI data was performed using the Configurable Pipeline for the Analysis of Connectomes (Sharad et al., 2014) (C-PAC, version 1.4.0) following a standard protocol:

- The anatomical images were skull-stripped, segmented into three tissue types (cerebrospinal fluid - CSF, white matter - WM, gray matter - GM) and registered to the MNI152NLin6ASym template provided by FSL (S. M. Smith et al., 2004). The individual gray matter masks were generated by keeping the voxels with a probability of over 25% in the gray matter probability maps and a temporal signal-to-noise ratio (tSNR) over the 15 percentiles of all tSNR values in the functional image. The gray matter group masks for every cohort were derived by averaging the gray matter probability maps across subjects and keeping the voxels with a gray matter probability over 25%. For the joint analysis, the gray matter group masks of every cohort were multiplied with each other. The voxel-based morphometry (VBM) analysis was performed using FSL-VBM (Douaud et al., 2007). The modulated grey matter images were smoothed with an isotropic Gaussian kernel (FWHM = 5 mm).
- Functional images were slice-time corrected, realigned, motion corrected, skull-stripped and registered to the anatomical images. Thereafter, the global mean intensity was normalized across the fMRI run, the nuisance signals were regressed-out (scanner drift, physiological noise and head motion signals) and the timeseries were bandpass-filtered (0.01 - 0.1 Hz). Next, the degree of functional connectivity (dFC) was calculated based on the voxel-wise Pearsońs correlation (*p* < 0.001 significance threshold) of the preprocessed timeseries of each voxel within the individual gray matter mask using the function 3dDegreeCentrality from AFNI (Craddock et al., 2016). Finally, the dFC-map was spatially smoothed (Gaussian filter, FWHM = 6 mm) and registered to the MNI152NLin6ASym 3 mm template through the anatomical image. The regression of the nuisance signals modeled the scanner drift using quadratic and linear detrending, whereas the physiological noise was modeled using the five principal components with the highest variance from the decomposition of white matter and cerebrospinal fluid voxel time series (CompCor) (Behzadi et al., 2007).
- The dynamic functional connectivity was calculated as the standard deviation over time of the degree of functional connectivity generated from sliding windows time series (Yu et al., 2015) (width = 40 s in steps of 20 s). The dynamic dFC map was spatially smoothed (Gaussian filter, FWHM = 6 mm) and registered to the MNI152NLin6ASym 3 mm template through the anatomical image.
- Diffusion-weighted images preprocessing and probabilistic tractography were performed using MRtrix3 (version 3.0.0) (Tournier et al., 2019), FSL (Jenkinson et al., 2012) and Advanced Normalization Tools (ANTs) (Avants et al., 2011), following the anatomically-constrained tractography pipeline (R. E. Smith et al., 2012). The preprocessing included denoising, eddy-current correction, motion correction (using FSL topup) and bias-field correction (using ANTs). The structural connectivity matrices were derived from the preprocessed images using single-tissue constrained spherical deconvolution probabilistic tractography (Tournier et al., 2007). Additionally, a spherical-deconvolution informed filtering was applied to the tractograms, constrained by the anatomical tissue masks and the HCP-MMP parcellation. Finally, the strength of the structural connectivity was derived from the communicability between each pair of brain regions (Crofts & Higham, 2009), capturing the communication capacity of direct and indirect connections.

#### PET

For the TUM cohorts, the first 45 minutes of the PET acquisition were reconstructed offline using the NiftyPET library (Markiewicz et al., 2018) based on the ordered subsets expectation maximization (OSEM) algorithm with 14 subsets, 4 iterations, and divided into 33 dynamic frames: 10 x 12 s, 8 x 30 s, 8 x 60 s, 2 x 180 s and 5 x 300 s. The attenuation-correction was based on the T1-derived pseudo-CT images (Burgos et al., 2014). For the VIE cohort, the first 40 minutes of the PET acquisition were reconstructed offline using the Siemens e7 reconstruction tool based on the OSEM algorithm with 21 subsets, 3 iterations, and divided into 30 dynamic frames: 24 x 5 s, 1 x 60s, 1 x 120 s, 1 x 300 s, 1 x 600 s and 2 x 1200 s post injection. The attenuation-correction was based on low-dose CT images from the participants (Sundar et al., 2018).

All reconstructed PET images were motion corrected and spatially smoothed (Gaussian filter, FWHM = 6 mm). The net uptake rate constant (Ki) was calculated using the Patlak plot model (Patlak & Blasberg, 1985) based on the last 5 frames of the preprocessed PET images (frames between 20 to 45 minutes) and the arterial input function derived from the preprocessed arterial blood samples. The cerebral metabolic rate of glucose (CMRglc) was calculated by multiplying the Ki map with the concentration of glucose in plasma of every participant, divided by a lumped constant of 0.65 (Wu, 2003). Finally, the CMRglc maps were partial volume corrected using the GM, WM and CSF masks derived from the T1 images using the iterative Yang method (Yang et al., 2017) and registered to the MNI152NLin6ASym 3 mm template through the anatomical image.

#### Arterial input function (AIF)

For the TUM cohorts, the blood time-activity curves (TAC) were preprocessed using the Turku PET Center command-line interface library TPCCLIB (Oikonen et al., n.d.) (version 0.7.5). First, the blood TAC were converted to plasma TAC using the b2plasma function, based on the reference FDG plasma/blood ratio function (Phelps et al., 1979) over time and the hematocrit value of each participant when measured, otherwise a reference value of 0.4/0.45 (female/male) was used (Chernecky & Berger, 2012). For the VIE cohort, the whole blood samples were centrifuged to measure the radioactivity in the plasma. For all cohorts, the plasma TAC was modeled using a sum of exponential functions, which was fitted to the middle of the reconstructed PET timeframes to derive the AIF (Feng et al., 1993). For four participants from the TUM.exp cohort without complete arterial sampling, the AIF was generated based on a population-based input function (PBIF) (Vriens et al., 2009) derived from all the participants from the TUM.

#### Population-based input function (PBIF)

The population-based arterial input function (AIF) was calculated as the average AIF across participants with arterial input sampling from our center (n = 16). The individual AIF were then normalized by the expected FDG concentration immediately after injection Cp*(0) (Equation 1) (Shiozaki et al., 2000):

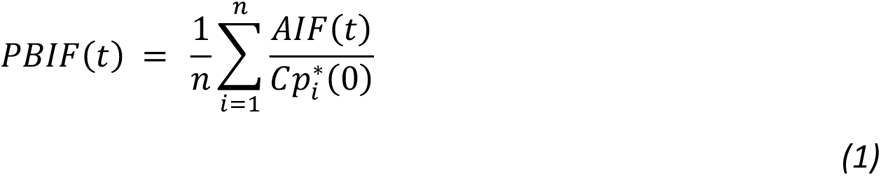

where *Cp _i_*(0)* is the initial plasma concentration of [18F]FDG, defined as the expected concentration directly after tracer injection, calculated by evaluating Equation *2* at t=0. Equation *2* models the plasma concentration of tracer as an exponential function during the period of intravascular and extravascular [18F]FDG equilibrium (Buxton, 2017), between 5 and 30 min after tracer injection (Sadato et al., 1998).

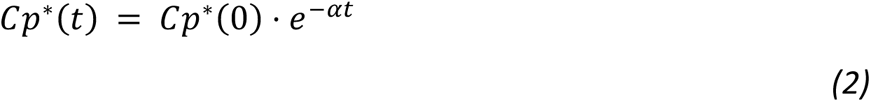

*α* was derived by fitting Equation *2* with the non-linear least squares optimization method from SciPy (Virtanen et al., 2020). For four participants of TUM.exp, who did not have arterial sampling due to technical errors, their individual AIFs (AIF_iPBIF_) were calculated based on the population-based input function (Equation *1*), scaled by the expected [18F]FDG concentration immediately after injection (Cp*(0)) (Equation *3*).

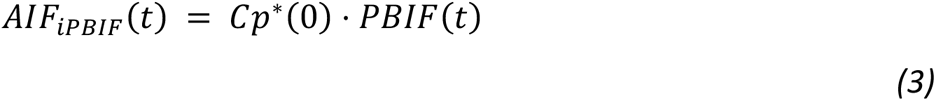

where *Cp*(0)* can be approximated as a function of the injected dose (iD), and the participant body weight (W) and height (H), when arterial sampling is not available (Equation *4*) (Shiozaki et al., 2000; Vriens et al., 2009).

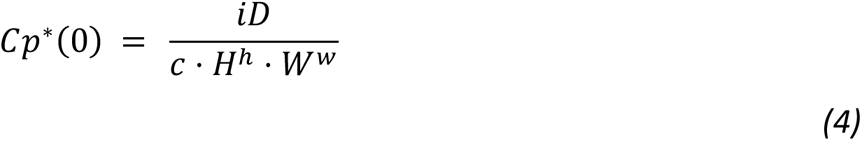

where *h*, *w* and *c* were derived by minimizing the coefficient of variation of 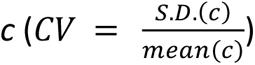 across the participants with arterial input sampling in our center, while varying independently the parameters *h* and *w* in the ranges of 0 to 2 and 0 to 1 in Equation *4*, respectively (Shiozaki et al., 2000).

#### Energetic costs of signaling

The relationship between brain energy metabolism (CMRglc) and the degree of brain-wide functional connectivity (dFC) was modelled using a linear regression model (Equation *5*) across cortical voxels within the GM mask.

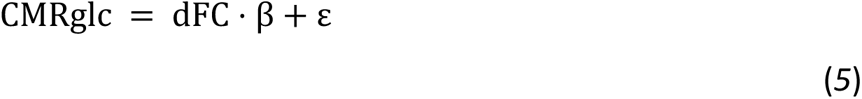

The energetic costs were defined as the residual after fitting the model in Equation *5*, representing the variation in CMRglc not explained by dFC. Positive energetic costs, located above the regression line, represent areas with a higher energy demand than expected for a given dFC, whereas negative energetic costs, located below the regression line, represent areas with a lower energy demand than expected for a given dFC.

### External data sources

Here, we integrated our group imaging data with 7 different external datasets about brain morphology, histology, transcriptomic data, molecular imaging, and comparative brain data. Each of the external datasets was provided in their own regional parcellation schemes and first converted to the HCP-MMP1.0 parcellation using the parcellation conversion tools provided by the Enigma-toolbox (Larivière et al., 2021).

1. *Allometric scaling between total brain metabolism and brain volume across species* (Karbowski, 2007) This dataset includes the CMRglc of unanaesthetised adult animals during resting conditions from ten different species, including humans, compiled from previous. The total glucose utilization rate was derived by multiplying the CMRglc with the correspondent brain volume.
2. *Cortical expansion from non-human primates to humans* (Wei et al., 2019, p. 201) This dataset provides a brain map of the cortical expansion from chimpanzees (Pan troglodytes) to humans (Homo sapiens) using surface-to-surface mapping of 3D cortical regions across both species based on in vivo T1-weighted MR-images of 29 chimpanzees and 30 humans.
3. *BigBrain Atlas* (Amunts et al., 2013) This dataset provides an ultra-high-resolution volumetric reconstruction of a postmortem, Merker-stained human brain from a 65-year-old. We used the version with 50 equivolumetric surfaces sampled between the pial and white matter surfaces provided by the BigBrain Warp toolbox (Paquola et al., 2021). First, the volume values were inverted to reflect the cellular density across cortical depth (Paquola et al., 2019), and then the skewness of each intensity profile was calculated to illustrate the distribution of cellular density between infra- and supra-granular layers (Zilles et al., 2002).
4. *Allen Human Brain Atlas (AHBA)* (Hawrylycz et al., 2012) This dataset provides microarray data collected across the entire cortex of six human donor brains. We used a preprocessed version of the data provided by the Enigma-toolbox (Larivière et al., 2021), based on the Abagen-toolbox (R. D. Markello et al., 2021). Preprocessing included the intensity-based filtering of microarray probes, the selection of a representative probe for each gene across hemispheres, the matching of microarray samples to brain parcels from the HCP-MMP1.0 parcellation, the normalization using the scaled robust sigmoid function across genes and samples, and the averaging within parcels and across donors of genes with a similarity across donors of at least 0.2, leaving a total of 8426 genes for further analysis. For the gene sets below, we calculated the region-wise mean and z-scored expression values for each gene.
5. *Genotype Tissue Expression database (GTEx v8)* (The GTEx Consortium, 2020) This dataset provides tissue-specific gene expression profiles collected from 54 non-diseased tissue sites from around 1000 participants. A list of 1588 genes that are expressed significantly more in the brain than compared to other organs was extracted based on a *p* < 0.05 (one-sided t-test, FDR corrected) and used as a background list for the gene ontology enrichment analysis (Wei et al., 2019).
6. *External PET neuroreceptors maps* (Hansen, 2022) This dataset compiles data from 28 different and previously published PET-imaging studies on the chemoarchitecture of the healthy human. It includes group-averaged volumetric PET maps of ligand occupancy from 19 unique receptors and transporters across 9 neurotransmitter and -modulator systems: dopamine, norepinephrine, serotonin, acetylcholine, glutamate, GABA, histamine, cannabinoid, and opioid. A subsequent analysis between transcriptomic and PET-imaging data included the following three PET ligands: the cholinergic nicotinic (acetylcholine) receptor alpha 4 beta 2 ([18 F]flubatine tracer), the opioid receptor mu 1 ([11 C]carfentanil tracer) and the serotonin 5-hydroxytryptamine receptor 4 ([11 C]SB207145 tracer).
7. *Neurosynth database* (Yarkoni et al., 2011) This dataset provides meta-analytic, statistical maps of brain activity related to 23 cognitive terms that are gathered from thousands of studies and ranging from sensorimotor to higher-order cognitive functions (Margulies et al., 2016). The statistically significant areas from the meta-analytic maps (z-score > 2.3) were used as masks to extract the distribution of the joint PLS score between the energetic costs and chemoarchitecture maps.

### Statistical analysis

#### Correlation analyses

The significance of the relationship between two variables was evaluated parametrically, based on the *p*-value associated to the Pearson’s correlation between them, and non-parametrically, based on the distribution of p-values derived from permuting the data 1000 times while preserving the spatial autocorrelation information of the target map (p_smash) using the Brainsmash toolbox (J. B. Burt et al., 2020, p. 202). To test for possible differences between cohorts and sex, these two variables were used as between-subject factors in independent one-way ANOVA analyses. The slope and correlation values derived from the linear relationship between CMRglc and dFC were used as the dependent variable. Additionally, the effect of age on the linear relationship between CMRglc and dFC was tested by calculating the Pearson’s correlation between the individual correlation values and age. The ANOVA and correlation analyses were performed using the python package Pingouin (version 0.3.9) (Vallat, 2018).

#### Analysis of spatial similarity between brain maps

The spatial similarity between the distribution of energetic costs of TUM.exp1 and those of the remaining cohorts was assessed based on a spatial Pearson’s correlation between them. The statistical significance of this correlation was evaluated by comparing it to a null distribution of correlation values derived from permuting the energetic costs of the TUM.exp1 map (1000 times) while preserving its spatial autocorrelation.

#### Statistical comparison between linear models

A one-way ANOVA was used to test for differences in the CMRglc variance explained by two linear models: a simple one with dFC as the only predictor variable, and a multiple linear model with dFC and other connectivity measures as additional predictors. The model comparison was performed in R Statistical Software (version 4.1.2) (R Core Team, 2022).

#### Allometric scaling

Allometry describes the scaling relationship of body parameters (Shingleton, 2010). Metabolic allometry of the brain models total brain size as a function of total glucose utilization rate according to the general allometric scaling law (Karbowski, 2007):

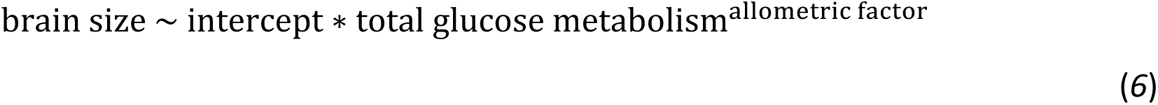

Equation *6* was fitted as a linear regression between log-log data of brain size and total glucose metabolism using the non-linear least squares optimization method provided by the python library SciPy (version 1.7.3) (Virtanen et al., 2020).

#### Group analysis of energetic costs

Brain regions with significantly deviating energetic costs were identified using voxel-wise one-sample t-tests of the individual parameter maps across all participants from all cohorts using the FSL randomize (Winkler et al., 2014) permutation-testing tool (*p* < 0.01, familywise error rate corrected, 5000 permutations).

#### Gene ontology enrichment (GOE) analysis

In this analysis we assessed the differentially expressed genes and their putative functions in brain regions of varying energetic costs. Across cortical regions, we correlated expression values of the 8426 AHBA genes with the average energetic costs map. Significantly correlated genes (*p* < 0.005, Benjamini-Hochberg FDR corrected) were subsequently used as an input to the GOE analysis and visualization tool GOrilla (version 03/06/2021) (Eden et al., 2009), and replicated using a different tool, Panther (version 17.0) (Mi et al., 2019). This analysis identifies gene ontology annotations for which the genes are significantly enriched using a minimal hypergeometric (mHG) *p*-value threshold of 10^-3^ (Eden et al., 2007, 2009), and corrected for multiple comparisons using a Benjamini-Hochberg FDR correction. As background, we used the brain specific gene set from the GTEx database (see External data sources).

#### Partial least square analysis (PLS)

The pyls python library (R. Markello, n.d.) was used to perform the PLS analysis between the z-scored map of energetic costs (334 ROIs x 30 participants) and the set of external PET-maps about the chemoarchitecture of the human brain (334 ROIs x 19 neurotransmitter receptors, see above). This analysis uses singular value decomposition to reveal the shared information between the two datasets represented as a set of orthogonal latent variables. The statistical significance of the latent variables was determined using permutation testing (5000 permutations), whereas bootstrap resampling (5000 times) was used to examine the contribution and reliability of the input features to each latent variable. The energetic costs score (surface representation in figure 4C) was calculated by projecting the energetic costs maps onto the first latent variable, whereas the receptor loadings (bar plot in figure 4C) are the Pearson correlation between the neuroreceptors map and the first energetic costs score.

