## Supplemental information for "An energy costly architecture of neuromodulators for human brain evolution and cognition"

This file includes:

Figs. S1 to S2
Tables S1 to S4

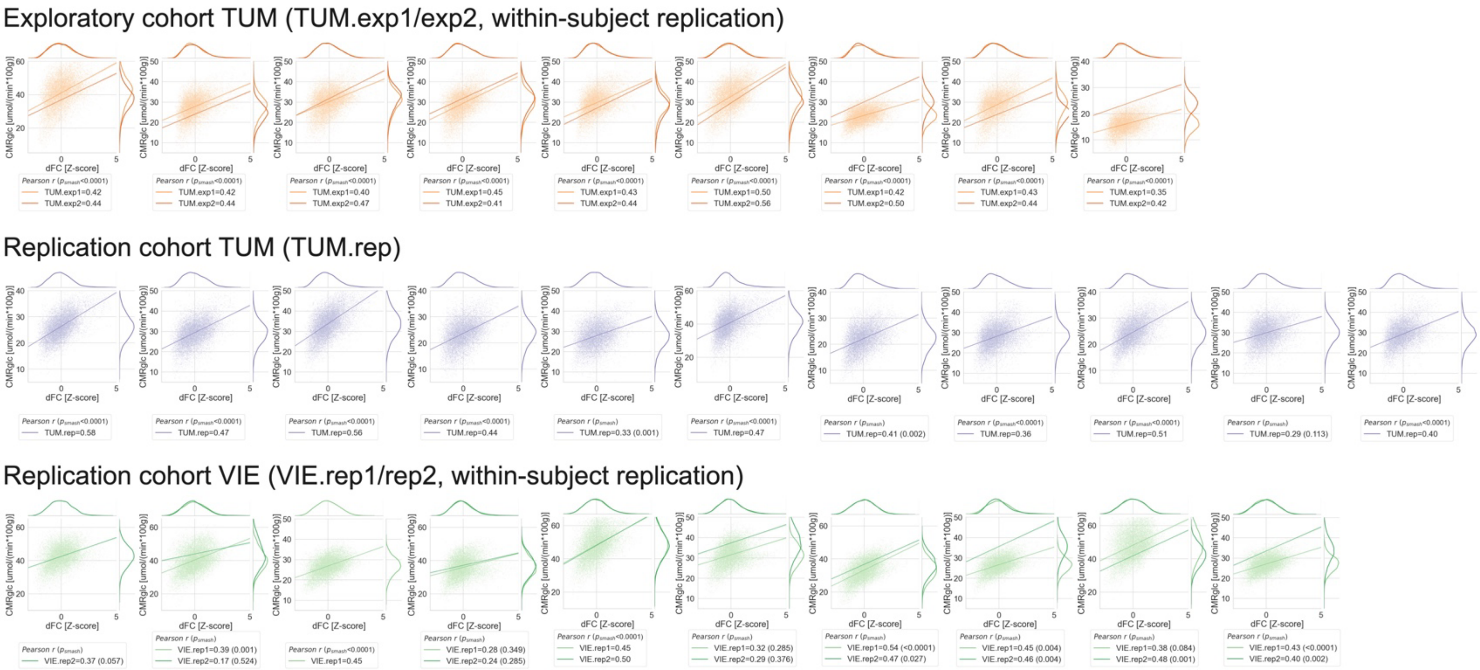


**Fig. S1: Glucose metabolism scales linearly with the degree of functional connectivity in individual subjects of all cohorts.** Significant Pearson's correlation between voxel-wise CMRglc and dFC for each dataset in all cohorts (*N* = 47, Pearson's *r*: mean = 0.42, s.d. = 0.08; all *p* < 0.1 * 10^-24, from individual correlation analyses; *p_smash* > 0.05 for *N* = 8 datasets; *N* ~ 11.000 voxels each). **Top:** TUM.exp1/exp2 (light/dark orange). **Middle:** TUM.rep (violet). **Bottom:** VIE.rep1/rep2 (light/dark green).

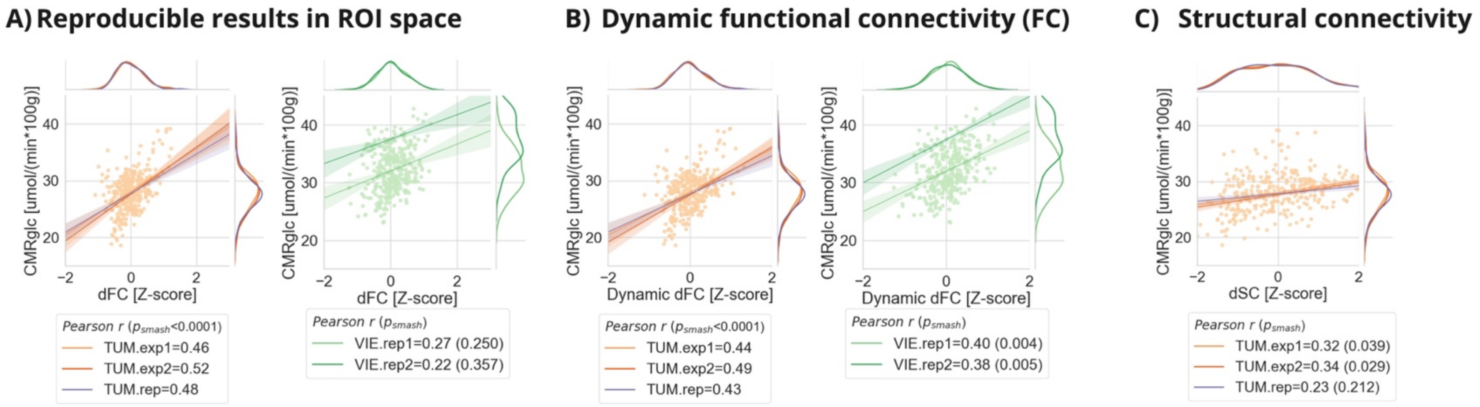


**Fig. S2: Stability of energy-connectivity scaling for different connectivity measures.** Glucose metabolism scales linearly with **A**, dFC, **B**, dynamic dFC, and **C**, structural connectivity averaged across individuals from each of the five cohorts using the MMP parcellation. (**A**) Significant linear relationship between region-wise CMRglc and dFC across all participants from the TUM cohorts complementing the voxel-wise results from the main text (all $p < 0.0001$; *p_smash* < 0.0001 for the TUM cohorts, correcting for spatial autocorrelation with permutation testing), explaining 22.5% of the total variance. (**B**) The significant linear relationship between region-wise CMRglc and dynamic dFC (all *p_smash* < 0.006, correcting for spatial autocorrelation with permutation testing) does not explain additional variance when added as an explanatory variable to the linear model between the CMRglc and the dFC (variance = 22.53%; $F(2,330) = 49.28$, $p = 0.29$, one-way ANOVA). (**C**) The significant linear relationship between region-wise CMRglc and degree of the structural connectivity (dSC) (*p_smash* < 0.05 for the TUM.exp cohorts and *p_smash* = 0.21 for the TUM.rep cohort; SC data was only available for the TUM cohorts) explains a significant additional variance when added as an explanatory variable to the linear model between the CMRglc and the dFC (variance = 28.78%; $F(2,330) = 68.08$, $p < 0.0001$, one-way ANOVA).

**Table S1**. Cell components significantly associated with co-expressed genes identified by the gene ontology analysis tool GOrilla.

| GO term | Description | p-value (FDR corrected) | Enrichment |
|---|---|---|---|
| GO:0044459 | plasma membrane part | 2.52E-5 | 1.44 |
| GO:0031226 | intrinsic component of plasma membrane | 2.2E-5 | 1.66 |
| GO:0005887 | integral component of plasma membrane | 4.96E-5 | 1.67 |
| GO:0044456 | synapse part | 2.57E-4 | 1.70 |
| GO:0098590 | plasma membrane region | 1.79E-3 | 1.59 |
| GO:0043005 | neuron projection | 3.63E-3 | 1.56 |
| GO:0097458 | neuron part | 3.12E-3 | 1.42 |
| GO:0030425 | dendrite | 1.02E-2 | 1.86 |
| GO:0005886 | plasma membrane | 9.54E-3 | 1.24 |
| GO:0045202 | synapse | 2.24E-2 | 1.55 |

**Table S2**. Molecular functions significantly associated with co-expressed genes identified by the gene ontology analysis tool GOrilla.

| **GO term** | **Description** | **p-value (FDR corrected)** | **Enrichment** |
|---|---|---|---|
| GO:0004930 | G protein-coupled receptor activity | 1.62E-2 | 2.22 |
| GO:0004888 | transmembrane signaling receptor activity | 1.06E-2 | 1.80 |
| GO:0022843 | voltage-gated cation channel activity | 1.3E-2 | 2.47 |
| GO:0038023 | signaling receptor activity | 1.37E-2 | 1.68 |
| GO:0001664 | G protein-coupled receptor binding | 1.51E-2 | 2.22 |
| GO:0060089 | molecular transducer activity | 4.26E-2 | 1.59 |
| GO:0005244 | voltage-gated ion channel activity | 4.84E-2 | 2.16 |

**Table S3**. Cell components significantly associated with co-expressed genes identified by the gene ontology replication analysis using the tool Panther.

| GO term | Description | p-value (FDR corrected) | Enrichment |
|---|---|---|---|
| GO:0043025 | neuronal cell body | 4.54E-02 | 2.42 |
| GO:0005911 | cell-cell junction | 4.54E-02 | 2.3 |
| GO:0120025 | plasma membrane bounded cell projection | 3.92E-02 | 1.47 |
| GO:0098978 | glutamatergic synapse | 3.69E-02 | 2.84 |
| GO:0030424 | axon | 2.84E-02 | 2.12 |
| GO:0042995 | cell projection | 2.50E-02 | 1.48 |
| GO:0044297 | cell body | 2.45E-02 | 2.35 |
| GO:0099240 | intrinsic component of synaptic membrane | 1.95E-02 | 6.2 |
| GO:0030425 | dendrite | 1.79E-02 | 2.31 |
| GO:0099699 | integral component of synaptic membrane | 1.78E-02 | 7.57 |
| GO:0098793 | presynapse | 1.42E-02 | 2.54 |
| GO:0097447 | dendritic tree | 1.11E-02 | 2.37 |
| GO:0070161 | anchoring junction | 1.08E-02 | 1.85 |
| GO:0045211 | postsynaptic membrane | 1.06E-02 | 5.02 |
| GO:0097060 | synaptic membrane | 9.04E-03 | 3.79 |
| GO:0098936 | intrinsic component of postsynaptic membrane | 7.82E-03 | 20.65 |
| GO:0043005 | neuron projection | 7.49E-03 | 1.84 |
| GO:0099055 | integral component of postsynaptic membrane | 7.33E-03 | 20.65 |
| GO:0044306 | neuron projection terminus | 5.89E-03 | 12.39 |
| GO:0098794 | postsynapse | 5.07E-03 | 2.71 |
| GO:0098590 | plasma membrane region | 3.22E-03 | 2.1 |
| GO:0036477 | somatodendritic compartment | 1.88E-03 | 2.45 |
| GO:0005886 | plasma membrane | 6.89E-04 | 1.41 |
| GO:0031226 | intrinsic component of plasma membrane | 1.76E-04 | 2.12 |
| GO:0071944 | cell periphery | 1.63E-04 | 1.42 |
| GO:0005887 | integral component of plasma membrane | 1.62E-04 | 2.17 |
| GO:0045202 | synapse | 3.25E-05 | 2.32 |
| GO:0030054 | cell junction | 2.78E-05 | 1.98 |

**Table S4**. Molecular functions significantly associated with co-expressed genes identified by the gene ontology replication analysis using the tool Panther.

| GO term | Description | p-value (FDR corrected) | Enrichment |
|---|---|---|---|
| GO:0022843 | voltage-gated cation channel activity | 1.08E-02 | 7.02 |
| GO:0038023 | signaling receptor activity | 2.01E-02 | 2.38 |
| GO:0060089 | molecular transducer activity | 2.35E-02 | 2.38 |
| GO:0001664 | G protein-coupled receptor binding | 2.45E-02 | 5.31 |
| GO:0005244 | voltage-gated ion channel activity | 3.85E-02 | 4.65 |
| GO:0022832 | voltage-gated channel activity | 4.33E-02 | 4.65 |
| GO:0004930 | G protein-coupled receptor activity | 4.53E-03 | 5.94 |
| GO:0004888 | transmembrane signaling receptor activity | 5.87E-03 | 2.95 |
